# Label-free and high-throughput removal of residual undifferentiated cells from iPSC-derived spinal-cord progenitor cells

**DOI:** 10.1101/2022.12.16.520699

**Authors:** Tan Dai Nguyen, Wai Hon Chooi, Hyungkook Jeon, Jiahui Chen, Daniel Ninio Roxby, Jerome Tan Zu Yao, Cheryl Yi-Pin Lee, Shi-Yan Ng, Sing Yian Chew, Jongyoon Han

## Abstract

The transplantation of spinal cord progenitor cells (SCPCs) derived from human induced pluripotent stem cells (iPSCs) has beneficial effects on treating spinal cord injury (SCI). However, the presence of residual undifferentiated iPSCs amongst their differentiated progeny poses a high risk as it can develop teratomas or other types of tumors post-transplantation. Despite the need to remove these residual undifferentiated iPSCs, no specific surface markers can identify them for subsequent removal. By profiling the size of SCPCs after a 10-day differentiation process, we found that the large-sized group contains significantly more cells expressing pluripotent markers. In this study, we employed a sized-based, label-free separation using an inertial microfluidic-based device to remove tumor-risk cells. The device can reduce the number of undifferentiated cells from an SCPC population with high throughput (*i*.*e*., > 3 million cells per minute) without affecting cell viability and functions. The sorted cells were verified with immunofluorescence staining, flow cytometry analysis, and colony culture assay. We demonstrated the capabilities of our technology to reduce the percentage of OCT4-positive cells. Our technology has great potential for the ‘downstream processing’ of cell manufacturing workflow, ensuring better quality and safety of transplanted cells.

## 1. Introduction

As a potential source to generate various cell types for cell therapy, the conversion of patient-derived somatic cells into induced pluripotent stem cells bypasses many ethical issues that are associated with the use of human embryonic stem cells (hESCs) (Takahashi and Yamanaka, 2006). Although these induced pluripotent stem cells (iPSCs) hold great potential in regenerative medicine, the tumorigenicity of iPSCs remains a significant hurdle for the safe therapeutic application of iPSC-derived differentiated cells (Hong et al., 2014; Yasuda et al., 2018; Yasuda and Sato, 2015; Zhang et al., 2011). Due to their unlimited self-renewal and pluripotency, the residual undifferentiated iPSCs can form teratomas in a dose-dependent manner (Residual Undifferentiated Cells During Differentiation of Induced Pluripotent Stem Cells In Vitro and In Vivo, 2012; Gropp et al., 2012; Hong *et al*., 2014). The potential tumorigenicity risk of the residual undifferentiated iPSCs depends on the number of undifferentiated cells, the features of the cell lines, (Yasuda *et al*., 2018) and culture adaptation (Lee et al., 2013a). Therefore, it is crucial to evaluate the tumorigenicity of iPSC-derived differentiated cells for each cell line and batch to ensure their safe therapeutic use.

Several removal methods of residual iPSCs or ESCs have been reported. These include cytotoxic antibodies (Selective Removal of Undifferentiated Human Embryonic Stem Cells Using Magnetic Activated Cell Sorting Followed by a Cytotoxic Antibody, 2012; Lim et al., 2011; Tan et al., 2009), fluorescence-activated cell sorting (FACS) (Ben-David et al., 2013; Fong et al., 2009; Tang et al., 2011), magnetic-activated cell sorting (MACS) (Haramoto et al., 2020; Onuma et al., 2013) and small molecules (Lee et al., 2013b; Lin et al., 2017b). In general, these methods rely heavily on cell surface markers, which lack sufficient specificity for pluripotent cells. On the other hand, alternative removal methods that utilize gene manipulation or chemical inhibitors (Ben-David and Benvenisty, 2014; Ben-David *et al*., 2013; Chung et al., 2006; Rong et al., 2012), may not be suitable for high-throughput cell therapy manufacturing.

Microfluidic spiral sorting is a cutting-edge technique used for the high-throughput separation of cells based on physical properties like size, shape, and density, making it ideal for label-free sorting (Bhagat et al., 2008; Kuntaegowdanahalli et al., 2009). This technology has proven effective in selecting specific subpopulations for stem cell therapy (Lee et al., 2014; Poon et al., 2015; Song et al., 2017; Yin et al., 2020). Given that the size of iPSCs decreases during differentiation (Lin et al., 2017a), we suggest that microfluidic spiral sorting could be a valuable tool for removing residual iPSCs from the iPSC-derived population based on size differences. By utilizing this technique, we can further improve the purity and quality of stem cell populations, paving the way for more efficient and effective stem cell therapies.

In this study, a high-throughput microfluidic MDDS sorter was developed to remove residual undifferentiated cells from populations of iPSC-derived spinal cord progenitor cells (SCPCs), which have great potential for spinal cord injury treatment (Dell’Anno et al., 2018; Kajikawa et al., 2020; Kumamaru et al., 2018). The sorter takes advantage of the differences in cell size by applying stronger drag forces on the larger undifferentiated cells. This enables the efficient separation of undifferentiated cells from the heterogeneous population of cells in microfluidic channels. The sorter uses label-free separation, is simple, cost-effective, and high-throughput, making it an ideal solution for the rapid and large-scale production of safer SCPCs. Here, following the safety quality control (QC) strategy of detecting OCT4-positive cells in iPSC-derived neural stem/progenitor cells (Sugai et al., 2021), the expression of OCT4 was analyzed using immunostaining, flow cytometry, and colony culture assay (Kuroda et al., 2012) to quantify the purity of our microfluidics-based sorted cells.

## 2. Results

### 2.1. SCPCs size profiling and its correlation with pluripotent biomarker

The differentiation efficiency of SCPCs was assessed on day 10 based on the expression of pluripotent marker OCT4 and neural progenitor marker SOX1, in comparison to iPSCs. High OCT4 expression and no SOX1 expression were observed in iPSCs. In contrast, while most SCPCs expressed SOX1, indicating successful differentiation into neural progenitors, some cells still expressed OCT4 (as highlighted by circles in **Figure S1 of SI**), suggesting incomplete differentiation or the presence of residual iPSCs.

To gain insight into the biophysical changes that occur during differentiation, we investigated changes in cell size by analyzing microscopy images and size histograms of suspended CLEC-iPSCs (Figure 1A) and day 10 CLEC-SCPCs (Figure 1B). Our results showed that SCPCs were more homogeneous, with approximately 74% of SCPCs ranging from 10 µm to 14 µm in diameter. On the other hand, the size distribution of iPSCs was slightly wider (*i*.*e*., approximately 76 % of iPSCs ranging from 15 µm to 21 µm in diameter). Furthermore, iPSCs were significantly larger than day 4 SCPCs (*p* < 0.001), with a mean difference of 4.5 µm. Interestingly, from day 4 to day 10, the nucleus size slightly decreased (*p*= 0.016). We confirmed a similar observation in BJ-iPSCs and BJ-SCPCs (**Figure S2A of SI**), indicating that the size decrease is not cell line-dependent.

**Figure 1.**
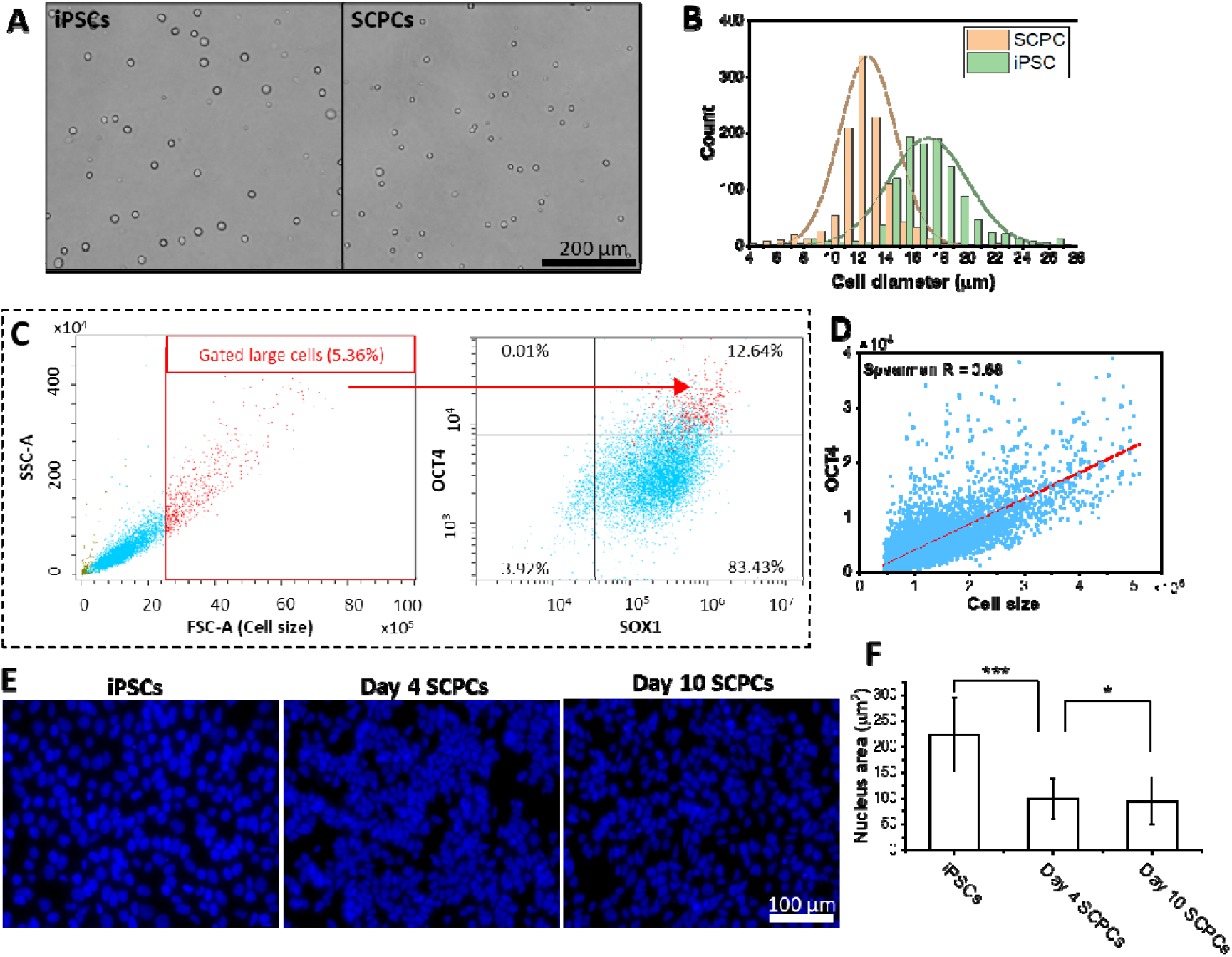
iPSC neural differentiation results in size and nucleus area reduction. (A) Microscopy images of suspended CLEC-iPSCs and CLEC-SCPCs. (B) Size profiling of the CLEC-iPSC and CLEC-SCPC using our customized MATLAB code. Cell size was significantly different, p < 0.001, Mann-Whitney test. N = 1326 in each group. (C) Flow cytometer analysis of the SCPCs. The large cells gated in the SSC-A/FSC-A plot are mostly OCT4-positive cells. The FSC intensity is proportional to the diameter of the cells in flow cytometry. (D) The OCT4 fluorescent intensity is correlated with the size of SCPCs (based on FSC intensity). Rank correlation by Spearman analysis (R = 0.68) indicates a moderate/good correlation between cell size and OCT4 level. The dashed line gated OCT4 positive and negative cells. (E) Immunostaining images with DAPI of the CLEC-iPSCs, and CLEC-SCPCs differentiated for 4 days and 10 days. The nucleus area of iPSCs was reduced during the differentiation. (F) Quantification of the nucleus area of the iPSCs, CLEC-SCPCs differentiated for 4 days and 10 days. The nucleus area was reduced significantly after 4 days (p < 0.001), and after 10 days (p = 0.016), Kruskal–Wallis test was followed by Dunn post-hoc test. N = 1200 for each group.

To verify our findings, we used flow cytometry, where the forward scatter intensity is approximately correlated to cell size and commonly used for gating cell types of different sizes. By gating out the FSC/SSC plot, we found that most of these cells fall into the OCT4^+^position of the OCT4/SOX1 plot (Figure 1C). Moreover, the intensity of OCT4 expression shows a good correlation to the cell size, as shown in Figure 1D, with Spearman’s rank correlation coefficient of 0.68. This result suggests that cell size is positively correlated with OCT4 expression, and the residual iPSCs retained their size morphology even after 10 days of differentiation.

Other than cell size, the nucleus area of the cells was significantly decreased during the differentiation. Figure 1E shows the immunostaining images with DAPI of the iPSCs, and SCPCs differentiated for 4 days and 10 days. By analyzing the nucleus area, we found that the iPSCs appeared to have a larger nucleus area than SCPCs on day 4 and day 10.

### 2.2. Sized-based separation of SCPCs and spiked iPSCs with the MDDS sorter

The size-based separation was achieved using a MDDS sorter, as illustrated in Figure 2**Figure 3**A. The sorter divided cells into four outlets: S2 (the smallest cells) to S5 (the largest cells), while cell fragments and debris were directed solely to outlet S1. This method was first demonstrated by performing a spiking experiment where a cell population containing 87% day 10 SCPCs spiked with 13% iPSCs was separated (Selective Removal of Undifferentiated Human Embryonic Stem Cells Using Magnetic Activated Cell Sorting Followed by a Cytotoxic Antibody, 2012; Ben-David *et al*., 2013; Haramoto *et al*., 2020; Wellmerling et al., 2020). The sorted cells were subsequently stained with pluripotency marker (OCT4), progenitor marker (SOX1), and spinal cord marker (HOXB4) and quantified through imaging with a fluorescent microscope to determine the cell composition (Figure 2B). Figure 2**Figure 3**C shows the size profiling and the percentage of OCT4 ^+^, SOX1 ^+^, and HOXB4^+^ cells in both unsorted and sorted groups. As iPSCs are typically larger than SCPCs, the S4 and S5 groups have a greater number of iPSCs stained with OCT4 (in red color). The S2 group, which comprises the smallest cells, exhibited the lowest OCT4 percentage at approximately 1%. Additionally, this group demonstrated a significantly higher percentage of HOXB4 expression and a stronger tendency towards SOX1 expression. This finding suggests that a larger proportion of these smaller cells may contain SCPCs. On the other hand, there was no significant difference observed in size and stained markers between the S4 and S5 groups. The difference in cell size between the S3 and S4 groups caused significant differences in the expression levels of OCT4, SOX1, and HOXB4. This indicates a clear distinction between small and large cells. Therefore, to reflect this observation, in the following sections of the paper, the cells will be separated into only two groups - Small and Large - instead of the initial four groups.

**Figure 2.**
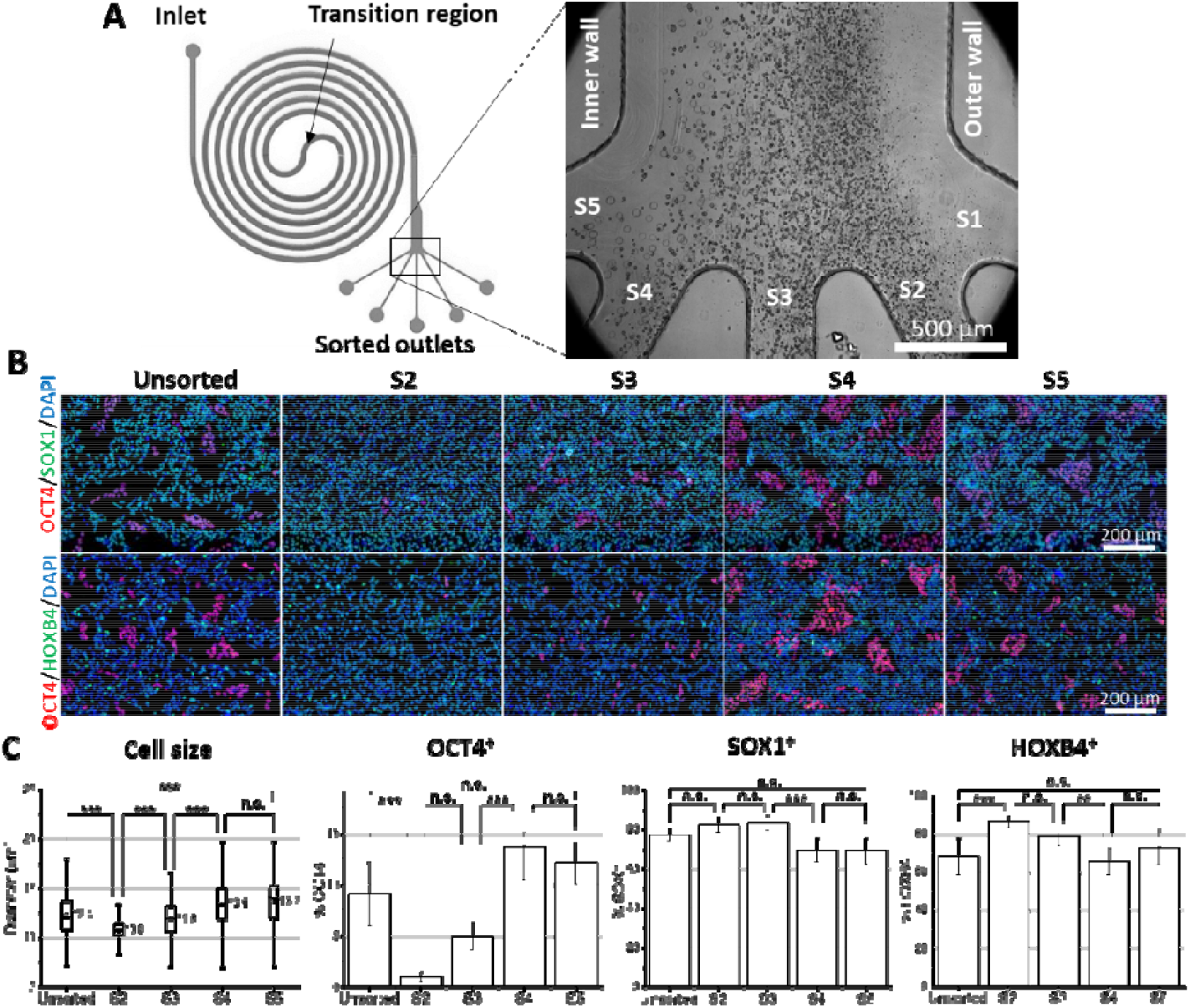
Sorting SCPCs and iPSCs using a microfluidic MDDS sorter. (A) Schematic of MDDS sorter (left) and representative bright-field image (right) showing cells were sorted into 5 outlets ranging from the smallest (S1) to the largest (S5). (B) SCPCs spiked with iPSCs were sorted using a MDDS sorter. The sorted cells were analyzed by immunofluorescence with OCT4 (red), SOX1 (green), HOXB4 (green), and DAPI (blue). (C) Quantification of cell size, OCT4, SOX1 and HOXB4 expression by unsorted group and sorted groups (i.e., S2 to S4). Cell size was significantly different, p < 0.001, Mann-Whitney test (n = 340). Significant difference in the expression of OCT4 and HOXB4 between the unsorted and S2 groups was observed. There was a significant difference in the expression of OCT4, SOX1, and HOXB4 between the S3 and S4 groups. Kruskal–Wallis test was followed by Dunn post-hoc test (n = 15).

**Figure 3.**
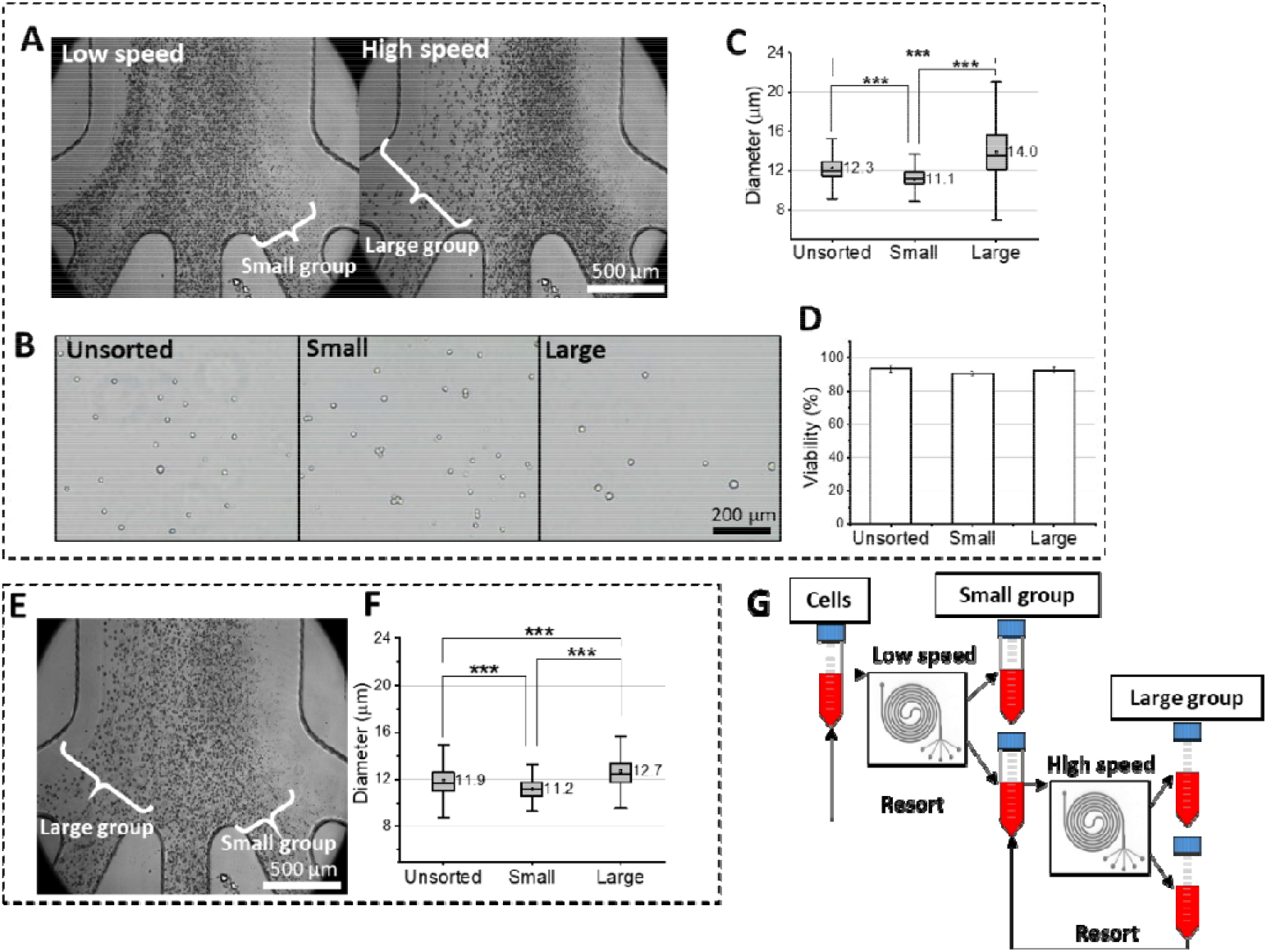
SCPCs sorting using microfluidic MDDS sorter. (A) Representative bright-field images showing cell separation under low (i.e., 2 ml/min) and high speed (i.e., 3 ml/min) operations or two-step operation, which allowed the collection of smaller and larger cells, respectively. (B) Representative microscopy images of Unsorted, Small and Large groups using two-step operation. (C) Box plot graph of cell size for the Small and Large groups (with a mean difference of 2.9 µm) compared to the Unsorted group. The cell size varies significantly among the groups (p < 0.001), as determined by the Kruskal-Wallis test, followed by a Dunn post hoc test (n = 324). (D) Viability of the Unsorted, Small and Large groups using two-step operation (n = 3). (E) Representative bright-field image showing cells sorted into Small (S2) and Large groups (S4 and S5) at a constant flow rate of 2 ml/min. (B, right). (F) box plot graph shows the size distribution of the Unsorted, Small and Large groups from E (mean difference of 1.5 µm). Cell size is significantly different among groups, p < 0.001, Kruskal-Wallis followed by a Dunn post hoc test (n = 324). (G) Schematic diagram depicting the recirculation strategy to re-sort cells that were not collected under low- and high-speed operations.

### 2.3. Optimization of MDDS sorter

We discovered an issue with overlapping cell sizes between sorted groups that resulted in a poor separation using the previous 5-outlet setting. To address this issue, we implemented a two-step operation of the MDDS sorter. By running the device at low speed (*i*.*e*., 2 ml/min), most cells focused on the inner wall except for small cells that were sorted to outlets S1 and S2 (*i*.*e*., Small group). Conversely, at higher speeds (*i*.*e*., 3 ml/min), most of the cells shifted to the outer wall, while very large cells were sorted to outlets S4 and S5 (*i*.*e*., Large group) (see Figure 3A). The size difference between the cells sorted into the Small and Large groups can be visually recognized from Figure 3B, while their sizes are shown in the box plot graph in Figure 3C. We also evaluated the viability of sorted cells and found that they remain highly viable, with no significant difference from the unsorted groups (Figure 3D). Compared to the constant-speed (i.e., 2 ml/min) sorting results in Figure 3E (microscopy image) and Figure 3F (box plot graph), the mean difference in cell size between the Small and Large groups increased from 1.5 µm to 2.9 µm when using the two-step operation. Furthermore, we implemented a recirculation strategy to re-sort uncollected cells to increase cell recovery for other analyses (see Figure 3G).

### 2.4. Immunofluorescent staining, colony culture assay, and flow cytometry analysis of sorted SCPCs: Reduced number of residual iPSCs

We employed the sorting of CLEC-SCPCs and characterized the removal efficiency of residual iPSCs. SCPCs were generated by following the standard 10-day differentiation protocol. The collected SCPCs were separated using a MDDS sorter with the recirculation strategy (resorted two times at low speed and three times at high speed). The three groups (Unsorted, Small, and Large) were stained for immunofluorescent imaging (Figure 4A). The cells that expressed OCT4 but not SOX1 (*i*.*e*., OCT4^+^SOX1^-^cells) were analyzed and shown in Figure 4B. The figure indicates that the Large group had more OCT4^+^SOX1^-^cells compared to the Unsorted group. In contrast, the Small group had slightly fewer OCT4^+^SOX1^-^cells. In addition, there was a trend indicating that the cells in the Large group had a larger nucleus area compared to the cells in the Unsorted and Small groups (Figure 4C). To assess whether the proliferation rate is correlated to cell size, EdU incorporation assays and Ki67 immunostaining were performed **(Figure S3)**. The results showed that iPSCs exhibited higher proliferation levels due to their capacity for unlimited self-renewal, while SCPCs displayed lower proliferation rates. Within the SCPCs population, smaller SCPCs showed a slower growth rate compared to larger SCPCs.

**Figure 4.**
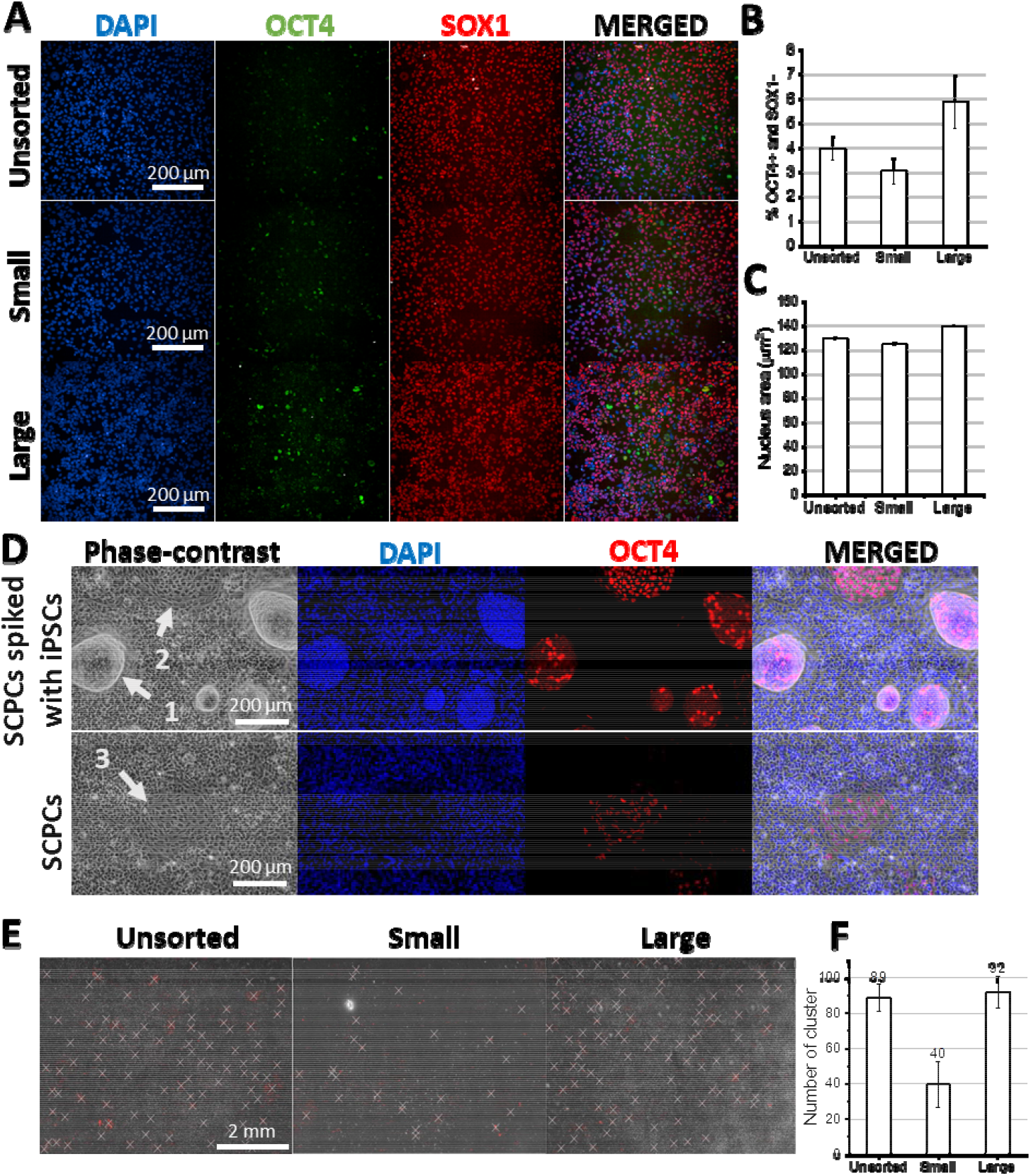
Immunostaining and colony culture assay of sorted CLEC-SCPCs. The SCPCs were separated into Small and Large groups by the recirculation strategy. (A) Immunofluorescence images of the Unsorted group and the sorted (Small and Large) groups with OCT4 (green), SOX1 (red), DAPI (blue), along with the merged images. (B) Percentage of OCT4^+^ SOX1^-^cells in the Unsorted, Small and Large groups (n = 3). (C) Comparison of cell nucleus area of the Unsorted, Small and Large groups (n = 3). (D) Colony culture assay. SCPCs and SCPCs spiked with iPSCs were cultured in an iPSC growth medium on Matrigel coating for 6 days before immunostaining and imaging. The SCPCs spiked with iPSCs were used as a positive control group to observe colony morphology formed from spiked iPSCs. Colonies were observed in phase contrast images of both SCPCs, and SCPCs spiked with iPSCs. Two types of colony morphology were observed: defined colony (with a sharp edge, compact cells pointed by number 1) and immature colony (with a defined edge, large nucleus pointed by number 2 for SCPCs spiked iPSCs group and number 3 for SCPCs groups). These colonies were verified by immunofluorescence images with OCT4 (red), DAPI (blue), and the merged image overlaid with phase contrast. (E) The map of identified colonies based on colocalization of phase contrast and OCT4 staining maps. The map consisted of 10 by 10 images, and identified colonies were marked with white “X”. (F) The number of quantified colonies in the Unsorted, Small and Large groups (n = 3).

Next, colony culture assay was performed by culturing SCPCs in a culture system that promotes the growth of iPSCs. Four separated cell populations, consisting of SCPCs spiked with 13% iPSCs (positive control group), the Unsorted, Small, and Large SCPCs groups were cultured in the 12-well plate. The cells were cultured in iPSC maintenance media for 6 days. Figure 4D shows the colony formation in the positive control group (SCPCs spiked iPSCs) and unsorted SCPCs group. The positive control group had well-defined colonies with inner packed cells and sharp edges (pointed by white arrow 1), which are the typical morphologies of iPSC colonies in culture. These colonies can also be identified in the DAPI staining, while the OCT4^+^cells are visible in the OCT4 staining. Besides the defined colonies, some colonies are in their early stages of colony formation with the large and elongated nuclei arranged in a circle (pointed by white arrow 2). For the unsorted SCPC group, similar immature colonies could also be found with the morphology of large nuclei arranged in circles (pointed by white arrow 3). Although immature colonies can be recognized by morphology, some colonies only express OCT4^+^marker (OCT4^+^aggregates) but do not have clear morphologies. Moreover, the distribution of OCT4^+^cells in the SCPC group was not always clustered in the form of colonies. Therefore, a density-based method of identifying OCT4 cells was developed to quantify the number of colonies formed among unsorted and sorted SCPCs (**Figure S4 in SI**). Quantified colonies in a map of 100 images (10 × 10 images) are shown in Figure 4E. Figure 4F presents a summary of the counted colonies obtained from the map of different groups. The figure indicates that the Unsorted and Large groups had a trend of a higher number of colonies compared to the Small group. These results indicate that the Small group has a much lower chance of forming colonies than the Unsorted and Large groups.

Figure 5A shows the dot plot with gated four-quadrant (SOX1 and OCT4) of unsorted and sorted cells from three batches of day 10 CLEC-SCPCs differentiated separately. To demonstrate that this strategy also works for other iPSC lines, we also utilized the same sorting approach to sort the day 10 SCPCs derived from BJ-iPSCs. As a starting point, both iPSC lines show a high level of pluripotency (> 98% expressed OCT4, **Figure S2B**). The OCT4 ^+^percentage of Unsorted, Small and Large groups for the SCPCs are presented in Figure 5B. The Large group exhibited a 2- to 5-fold increase in OCT4^+^ cells compared to the Unsorted group across different batches and cell lines. Similar to immunostaining results, some of the cells among OCT4 SCPCs also express SOX1. The percentage of OCT4^+^ SOX1^-^ is presented in Figure 5C, where batch 3 of CLEC-SCPCs and BJ-SCPCs show a significantly higher number of OCT4^+^ SOX1^-^ cells. This can be attributed to the higher OCT4^+^ cells in these groups. For both OCT4 and OCT4 SOX1 quantifications, all Small groups showed a lower percentage than the Unsorted group. Figure 5D provides an estimate of the removal efficiency of OCT4^+^ and OCT4^+^ SOX1^-^cells by comparing their cell count in the Large group to the initial cell number, which represents the number of these cells before they were sorted. Across different batches and cell lines, the results show that more than 40 % of OCT4+ cells and 30 % of OCT4^+^ SOX1^-^ cells can be removed, except for batch 1 of CLEC-SCPC, which had a high removal rate of OCT4^+^ cells but a low removal rate of OCT4^+^ SOX1^-^ cells. On the other hand, sorting of BJ-SCPCs showed an impressive elimination of 98 % of OCT4^+^ SOX1^-^ cells.

**Figure 5.**
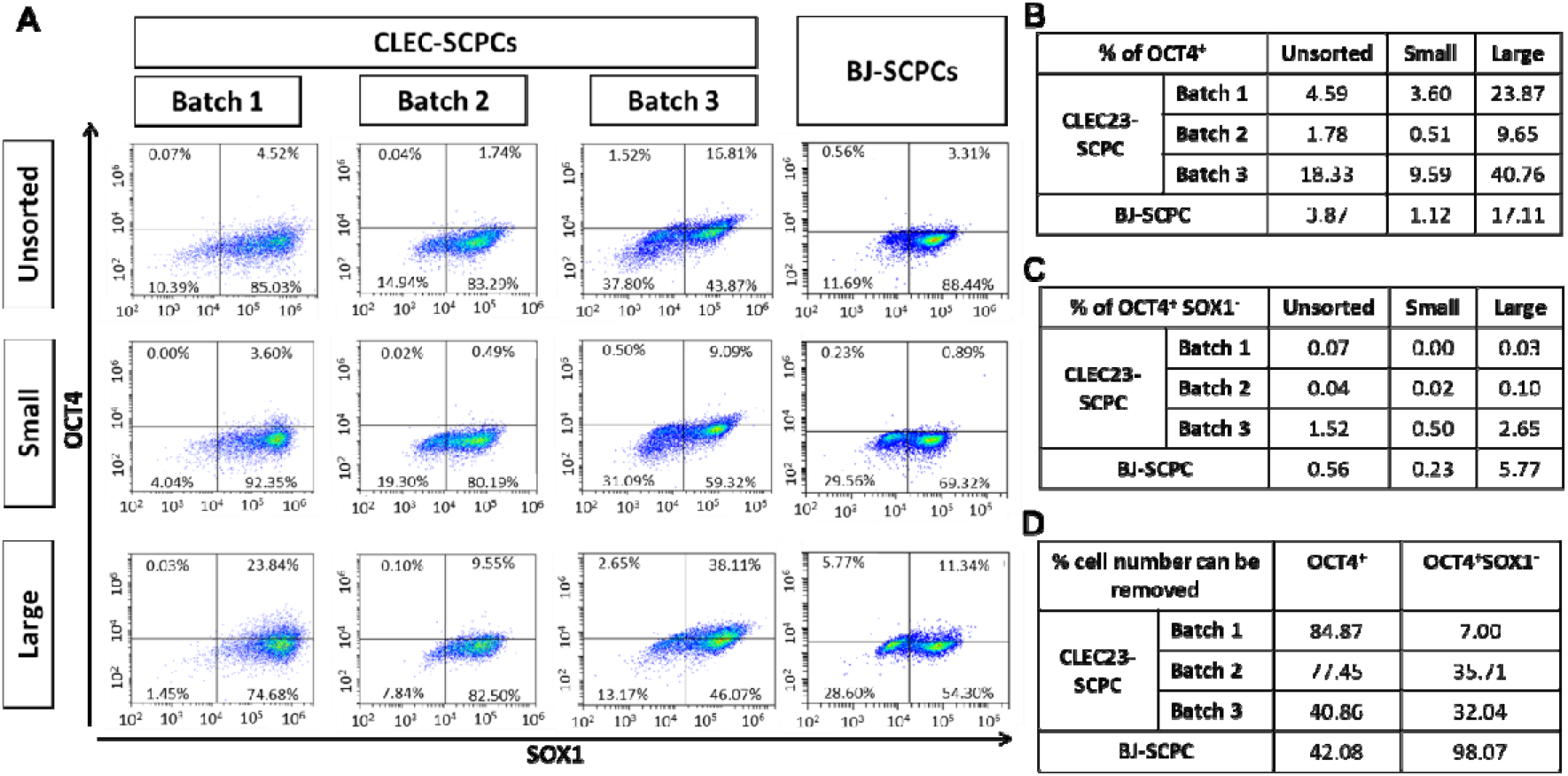
Flow cytometry assay shows that the percentage of OCT4^+^cells is consistently lower in the sorted Small group regardless of cell culture batches and cell lines. (A) Flow cytometry analysis of the Unsorted group and the Small and Large groups from three batches of day 10 CLEC-SCPCs generated separately and day 10 BJ-SCPC. (B) and (C), the percentage of OCT4^+^ and OCT4^+^ SOX1^-^ cells, respectively, of the Unsorted, Small and Large groups. (D) The percentage of OCT4^+^ and OCT4^+^ SOX1^-^ cell in the Large group compared to the unsorted cells.

## 3. Discussion

Due to its self-renewal and tumorigenic traits, residual iPSCs pose a high risk of forming tumors after transplantation. Much evidence in the past decade has demonstrated tumor formation from residual iPSCs in preclinical studies. Therefore, the removal of residual iPSCs is critical for safe cell therapy. Our study suggests that the large-sized cell group possesses more OCT4^+^ cells. These cell characteristics indicate their similarity to residual iPSCs and imply that residual undifferentiated cells tend to retain their biophysical properties, such as size, deformability, or shape, during differentiation. By exploiting the size difference between undifferentiated and differentiated cells, we developed a microfluidic size-based sorter for the removal of residual iPSCs in a label-free and high-throughput manner. Sorted cells were examined with immunofluorescence staining, a culture system for colony formation, and flow cytometry. Through both immunofluorescence staining and flow cytometry analysis, we found that a subset of OCT4^+^ cells also expressed the SOX1 marker, suggesting that these cells were in the transition stage of differentiation into SCPCs. The minority population that expresses OCT4 ^+^ (a pluripotent marker) but not SOX1 (a differentiation marker) would pose a higher risk of forming tumors. Additionally, we observed a significantly higher percentage of OCT4^+^ SOX1^-^ cells in the Large group than in the Small and Unsorted groups. However, immunostaining analysis of OCT4^+^ cells can be only qualitative, not quantitative, due to weak expression and errors in image-based quantification. On the other hand, flow cytometry is a more common method for the detection of residual undifferentiated cells due to high-speed analysis of a large number of cells (*e*.*g*., 100000 cells/second) and the capability of identifying even the minor populations (*i*.*e*., OCT4^+^ cell population) (Residual Undifferentiated Cells During Differentiation of Induced Pluripotent Stem Cells In Vitro and In Vivo, 2012; Fong *et al*., 2009; Kuroda *et al*., 2012). In this study, forward scatter intensity in the flow cytometer can also give us information about the cell size and its relation to other parameters.

Other than cell size, the nucleus area of the cells was significantly decreased during the differentiation. By analyzing the nucleus morphology, we found that iPSCs appeared to have a larger nucleus area than SCPCs on day 4 and day 10 (Figure 1E). This result is consistent with Figure 4C, where Large group cells had a larger nucleus area than Small group cells. The nucleus area could be a good quality attribute for differentiating residual iPSCs in SCPCs or other progenitor cells. This and all the other analyses collectively corroborate our central hypothesis that size-based sorting of SCPCs can eliminate undifferentiated cells, reducing risk.

It has previously been suggested that the slow-dividing neural progenitor cells in culture might be better for clinical applications (Furutachi et al., 2015). Meanwhile, hyperproliferative cells (*i*.*e*., possibly tumorigenic cells) may be caused by the genetic and/or epigenetic transformation of neural progenitor cells, not by the contamination of undifferentiated iPSCs (Iida et al., 2017; Sugai et al., 2016). We currently do not have a detailed understanding of the molecular pathways or mechanisms behind the cell size difference correlated with the phenotype shift we observe in iPSCs and their differentiated progenitors. Regardless of the source of the fast-growing residual cells, the high proliferation of the Large group could be associated with higher tumorigenic potential and, therefore, should be removed to reduce such a risk. At the same time, as suggested by some earlier studies, subpopulations with different sizes could represent cells with different phenotypes in therapeutic cells, which we previously addressed with the same cell sorting technology we demonstrated in this work (Lee *et al*., 2014; Poon *et al*., 2015).

Quality control of stem cells should be mainly discussed in terms of suitability for manufacturing the final products, especially the cell products derived from iPSCs (Jha et al., 2021). Our MDDS sorter will contribute to the high throughput removal of residual iPSC contaminations. Previously, we demonstrated the sorting throughput of 1L/min corresponding to ∼500 million cells/min by using a multiplexed plastic spiral unit containing 100 MDDS sorters (Jeon et al., 2022b). Therefore, scale-up to manufacturing scale cell production processes is feasible and straightforward.

In order to demonstrate general applicability, we tested our idea on two different iPSC cell lines and multiple batches. Based on the calculation of OCT4^+^ SOX1^-^ percentage from flow cytometry assay and sorted cell number, we can eliminate approximately 32.4% of OCT4^+^ SOX1^-^ cells from day 10 CLEC-SCPC population and approximately 90.6 % OCT4 SOX1 cells from day 10 BJ-SCPC population by removing the Large group (Figure 5D). While the method works for different iPSC lines, the differences in size distribution and differentiation efficiency also validated that the differentiation might not be fully efficient and can be cell origin dependent (Hargus et al., 2014; Hu et al., 2010; Strano et al., 2020). We observe significant variations in the relative amount of undifferentiated residual cells, not only for cells from different cell lines but also in different batches from the same cell lines (Cahan and Daley, 2013; Hayashi et al., 2019). The sources of such variation are not understood currently and could result from cell source, donor, culture conditions, quality of raw materials used in culture, and operator, to name a few. While it may be feasible to minimize or eliminate such variation by controlling cell culture and production conditions accurately (Rohani et al., 2018; Rosati et al., 2018), the availability of an easy-to-use ‘downstream’ purification for cell products would still be highly valuable and desirable in large-scale cell manufacturing.

In our inertial microfluidic cell separation, the estimated removal efficiency varies from 40-90%, depending on the cell lines and batches. A complete baseline separation between SCPCs and residual cells is not likely to be possible because of the overlap in size between fully differentiated SCPCs and undifferentiated residual cells. Further optimization of the sorting devices, combined with multiple rounds of removal separation, could lead to a satisfactory and acceptable level of residual cell removal, which would still be quantifiable in terms of enhancing the safety of the cells. At the same time, the cost of the device (injection-molded, mass-produced plastic chips) is expected to be minimal, and there are no reagents needed for the removal operation. Therefore, our device would provide a highly economic yet tangible benefit to any iPSC-derived cell products for regenerative medicine. In contrast, conventional FACS sorting would be not only limited in terms of the processing rates and throughput (if based on size sorting by scattering) but also prohibitively expensive if the sorting is based on specific cell-surface marker labels.

In conclusion, we utilized the size differences between the undifferentiated and differentiated cells to enrich the SCPCs population while reducing the percentage of iPSCs. The technology is label-free, non-contact, and has high throughput without affecting cell viability and functions. While we demonstrate our technique using a specific cell manufacturing scenario (transplantable SCPCs derived from iPSC cell line), we argue that the methodologies demonstrated here will find broad use cases in many clinical indications where cell purity and quality become an important issue. Further development of this technology will be promising for the cell manufacturing workflow, contributing to better quality and safety of the transplanted cells by reducing the number of potentially tumor-forming cells.

## 4. Experimental procedures

### 4.1. Resource availability

Corresponding author: Further information and requests for resources and reagents should be directed to and will be fulfilled by the corresponding authors: Jongyoon Han and Sing Yian Chew.

#### Materials availability

This study did not generate new unique reagents.

#### Data and code availability

All codes that enable the main steps of the analysis and data are available from the corresponding authors under request.

### 4.2. SCPC differentiation from human iPSCs

The iPSC lines were maintained in StemMACS iPS-Brew XF cell culture medium on Matrigel Matrix coating (Corning, USA). Healthy umbilical cord lining-derived iPSCs (CLEC23) (provided by Dr Kah-Leong Lim) are hypothesized to have the immune-privileged properties of the CLECs (Lim et al., 2020; Saleh and Reza, 2017). BJ-iPSCs were derived from BJ-fibroblasts using modified mRNA, which represent the conventional fibroblast-derived iPSCs and have been used to generate spinal motor neurons (Hor et al., 2021; Ng et al., 2015). The iPSCs were routinely passaged every 5-7 days using ReLeSR™ Passaging Reagent (Stemcell Technologies, Canada). The iPSCs were then differentiated into SCPCs based on a modified protocol to generate spinal motor neurons (Hor et al., 2018; Winanto et al., 2020). Briefly, when the iPSCs were 70-80% confluent, the iPSCs were dissociated into single-cell suspension using Accutase (Nacalai Tesque Inc., Japan). The dissociated iPSCs were then counted and reseeded at a density of 800,000 cells per well in a 6-well plate in a neural induction medium supplemented with ROCK inhibitor Y-27632 (5 µM). The neural induction medium consisted of DMEM/F12 (50%, Biological Industries, Israel), neural medium (50 %), NeuroBrew-21 (1X), N2 (1X), Non-Essential Amino Acid (1X, Thermo Fisher Scientific, USA), Glutamax (0.5X, Thermo Fisher Scientific), LDN-193189 (0.5 µM), and CHIR-99021 (4.25 µM). On day 3, retinoic acid (RA, 10 µM, Sigma-Aldrich, USA) was added to the medium. On day 4, the cells were reseeded at a lower density (2 million cells per 100 mm dish) in the same medium and maintained with daily medium change until day 10. On day 10, the cells were characterized and termed as SCPCs, which were then frozen or used for experiments. Unless specified, all other culture components were purchased from Miltenyi Biotech (Germany).

### 4.3. Microfluidic MDDS sorter design and fabrication

The polydimethylsiloxane (PDMS) MDDS sorter device was fabricated via the standard soft-lithographic technique. 3D CAD software (SolidWorks 2020, USA) was used to design an aluminum mold for making the PDMS MDDS sorter replica, and the aluminum mold was fabricated via a micro-milling process (Whits Technologies, Singapore). Here, we used the multi-dimensional double spiral (MDDS) inertial microfluidic device as the MDDS sorter (Jeon et al., 2022a; Jeon et al., 2020; Jeon et al., 2021), and its optimal channel dimensions and channel configuration were determined based on the observation of trajectories of particles with various sizes (7-30 μm) in the devices with different channel dimensions. The MDDS device is comprised of two sequentially interconnected spiral channels with distinct dimensions. The design of the first spiral channel incorporates a comparatively smaller dimension to generate an increased inertial lift force, subsequently directing all target particles or cells towards the inner wall side of the channel. In contrast, the second spiral channel features an enlarged dimension, allowing particles to attain disparate equilibrium positions as dictated by the balance between inertial lift and Dean drag forces, ultimately facilitating particle separation based on size (Jeon *et al*., 2022a). Through empirical selection, an optimal device was identified, in which the first spiral channel exhibits a rectangular cross-section with an 800 μm width, 100 μm height, and three loops, while the second channel demonstrates a trapezoidal cross-section, encompassing an 800 μm width, heights of 120 and 180 μm for the inner- and outer-wall sides respectively, and three loops.

A 10:1 mixture of PDMS base and curing agent (Sylgard 184, Dow Corning, Inc., USA) was used to make the PDMS replica. After curing on the hot plate for 10 min at 80 °C, the PDMS replica was bonded to a glass substrate using a plasma machine (Femto Science, South Korea).

### 4.4. SCPCs sorting using MDDS sorter

Prior to sorting, the MDDS sorter was incubated with 70% ethanol for at least 30 min for sterilization. It was then rinsed with 1X Phosphate-buffered saline (PBS) and medium. The cells were loaded into a 20 ml syringe and injected into the device at a flow rate of 2 ml/min for low-speed mode and 3 ml/min for high-speed mode using a syringe pump (PHD ULTRA Syringe Pumps, Harvard Apparatus, USA). To implement the recirculation strategy, a dual check valve (Quosina, USA) was used to retract the sorted cells into the input syringe by withdrawing them at a flow rate of 10 ml/min from the outlet reservoir using the syringe pump.

### 4.5. Immunofluorescent staining

The sorted cells were seeded at a density of 80,000 cells per well in a 96-well plate. After the cells were attached to the wells overnight, the cells were fixed in 4% paraformaldehyde (PFA, Santa Cruz Biotechnology, USA) for 15 min at room temperature. Thereafter, the cells were permeabilized in 0.1% Triton X-100 (Sigma-Aldrich, USA) for 15 min, followed by 1 h incubation in the blocking buffer (5% Fetal Bovine Serum (FBS, Gibco, USA) and 1% Bovine Serum Albumin (BSA, Sigma-Aldrich, USA) in PBS (Gibco, USA)) at room temperature. The cells were then incubated in primary antibodies diluted in the blocking buffer at 4 °C overnight. The primary antibodies used were: OCT4 (1:500, sc-5279, Santa Cruz Biotechnology), SOX1 (1:500, 4194S, Cell Signaling Technology, USA), and HOXB4 (1:200, ab133521, Abcam, UK). On the next day, the cells were washed with PBS and incubated in respective secondary antibodies (donkey anti-mouse AF 488/donkey anti-rabbit AF 568, 1:1000, Thermo Fisher Scientific) and DAPI diluted in blocking buffer (1:1000, Thermo Fisher Scientific) for 1 h at room temperature. The cells were then washed with PBS before imaging. Imaging was performed on a Leica DMi8 microscope or Opera Phenix High-Content Screening System using 10× or 20× objectives.

### 4.6. Flow cytometry analysis

Cells were collected in 15 ml centrifuge tubes and adjusted to 2 × 10^6^ cells/tube. They were then fixed in 4% PFA and incubated at room temperature for 15 min. Thereafter, the fixed cells were pelleted by centrifugation at 3000 rpm for 5 min and washed once with PBS. Next, cells were stained by incubation at room temperature for at least 2 h with a primary antibody in perm/block buffer (0.5% saponin (Sigma-Aldrich, USA), 1% BSA). The primary antibodies used were: OCT4 (1:500, sc-5279, Santa Cruz Biotechnology, USA) and SOX1 (1:500, 4194S, Cell Signaling Technology, USA). Cells were centrifuged and washed twice with PBS. They were then stained by incubation in the dark at room temperature for at least 45 min with a secondary antibody in perm/block buffer. The secondary antibodies used were donkey AlexaFluor488-conjugated anti-rabbit IgGs (A-21206, Thermo Fisher Scientific) and donkey AlexaFluor555-conjugated anti-Mouse IgGs (A-31570, Thermo Fisher Scientific). After that, cells were washed with PBS and resuspended in PBS. Before running with the Cytoflex flow cytometer (Beckman Coulter, USA), cells were filtered through a 45 μm cell strainer to remove cell clumps and kept on ice. Compensation was performed and applied to the analysis. Due to the relatively weak expression level of the OCT4 marker, the percentage of OCT4^+^ cells was obtained using fluorescence minus one (FMO) control (i.e., FMO-DAPI-SOX1 control).

### 4.7. EdU-based proliferation assay

The proliferation of SCPC and hiPSC was measured with the Click-iT EdU flow cytometry assay kit (Invitrogen, USA). EdU (5-ethynyl-2’-deoxyuridine) is a nucleoside analog to thymidine and is incorporated into DNA during active DNA synthesis. The EdU was added (10 μM) to the culture medium before seeding for incubation. After ∼24 h of incubation, cells were harvested and rinsed with PBS containing 1% BSA. Thereafter, the cells were fixed with Click-iT fixative for 15 min in the dark at room temperature and rinsed in PBS with 1% BSA. The cells were then permeabilized with 1X Click-iT® saponin-based permeabilization for 15 min at room temperature in the dark. After that, they were incubated for 30 min at room temperature in the dark with a Click-iT reaction cocktail. Cells were washed and resuspended with 1X Click-iT® saponin-based permeabilization and wash reagent before being analyzed by Cytoflex flow cytometer (Beckman Coulter, USA).

### 4.8. Colony culture assay

Cells were seeded in a 12-well plate at an initial density of 17,000 cells per cm^2^ based. They were cultured in hiPSC medium for 6 days with daily change of medium. Cells were fixed and stained with DAPI and OCT4 antibodies before imaging under a Leica DMi8 microscope.

### 4.9. Statistical analysis

The data were subjected to normality and homogeneity of variances testing using the Shapiro-Wilk and Levene’s tests, respectively. In cases where the data met the assumptions of normality and homogeneity, a one-way ANOVA followed by the Turkey post-hoc test was conducted to compare three or more groups. Otherwise, a nonparametric Kruskal–Wallis test followed by the Dunn post-hoc test was used. The results were presented as mean ± standard deviation (SD), with statistical significance indicated by ns (not significant), * (p ≤ 0.05), ** (p ≤ 0.01), or *** (p ≤ 0.001). Origin software (OriginLab Corporation, USA) was performed for the statistical analysis.

## Supporting information

C:\Users\nguyentd\OneDrive\Sorting paper\Submission

## 5. Author contributions

J. H, T.D.N, conceptualized the study and designed the study. H.J designed and fabricated the MDDS sorter. T.D.N, W.H.C, J.C, D.N.R, J.T.Z.Y, C.Y.P.L performed experiments. T.D.N, W.H.C analyzed data. S.Y.C, S.Y.N provided scientific advice and materials; J. H and S.Y.C conceived the project and supervised the research. T.D.N wrote the draft of the manuscript with input from W.H.C, H.J, J. H, S.Y.C. All of the authors read, edited, and approved the final version of the manuscript.

## 6. Acknowledgments

This research was supported by the National Research Foundation, Prime Minister’s Office, Singapore under its Campus for Research Excellence and Technological Enterprise (CREATE) programme (IntraCREATE grant award number: NRF2019-THE002-0001) and Singapore MIT Alliance for Research and Technology (SMART): Critical Analytics for Manufacturing Personalised-Medicine (CAMP) Inter-Disciplinary Research Group. We thank Prof. Kah Leong Lim (Lee Kong Chian School of Medicine, Nanyang Technological University) for providing the umbilical cord lining-derived iPSCs (CLEC23).

