## Supplementary material for "Label-free and high-throughput removal of residual undifferentiated cells from iPSC-derived spinal-cord progenitor cells": C:\Users\nguyentd\OneDrive\Sorting paper\Submission

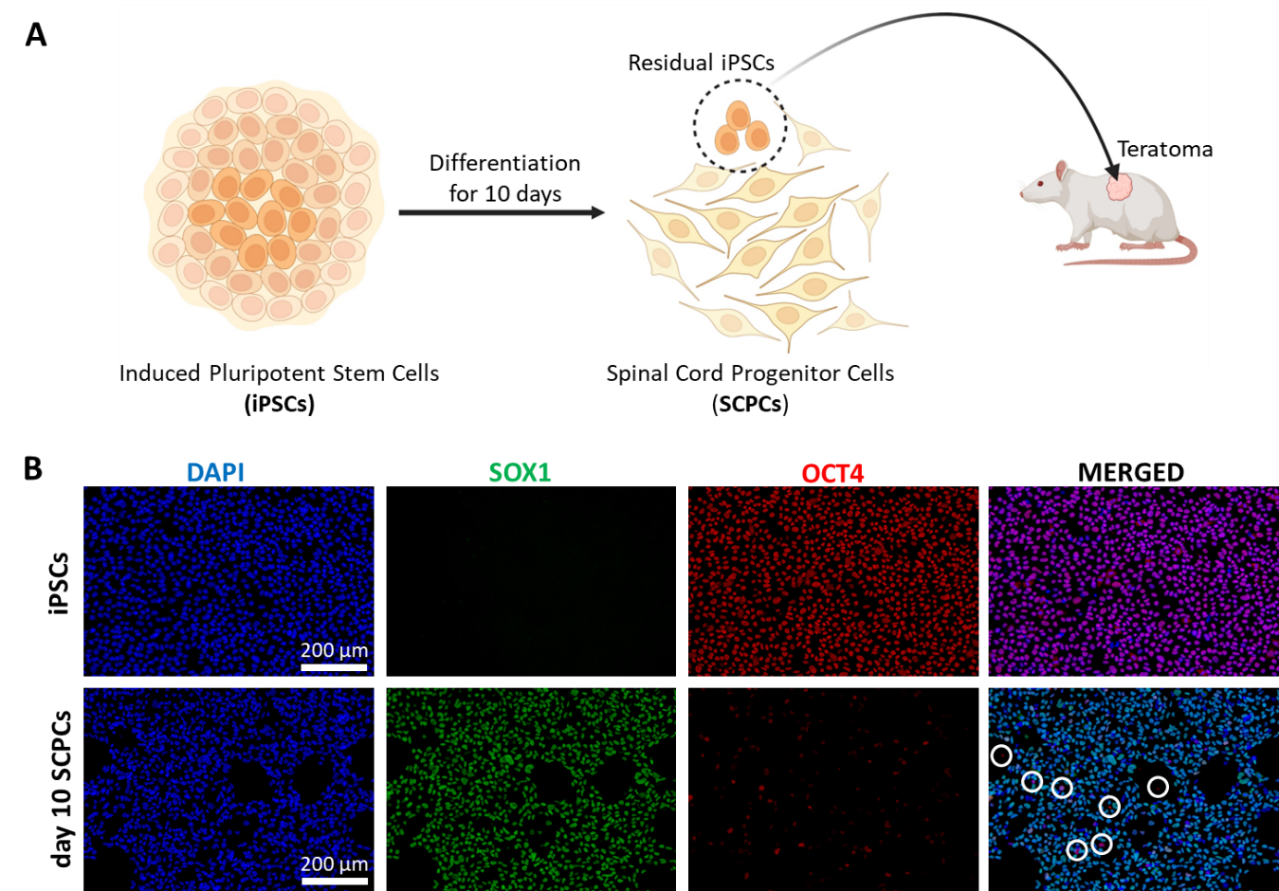

Figure S1 Tumor risk of residual iPSCs in the Spinal Cord Progenitor Cells. (A) iPSCs are differentiated for 10 days to become SCPCs. However, some residual iPSCs in the SCPC population pose a risk of forming tumors after transplantation (B) Immunofluorescent staining of iPSC and CLEC-SCPCs at day 10 of differentiation with OCT4 (red, pluripotent marker), SOX1 (green, progenitor marker), and DAPI (blue). In the merged image, some SCPCs (marked by the circles) still express OCT4 but not SOX1.

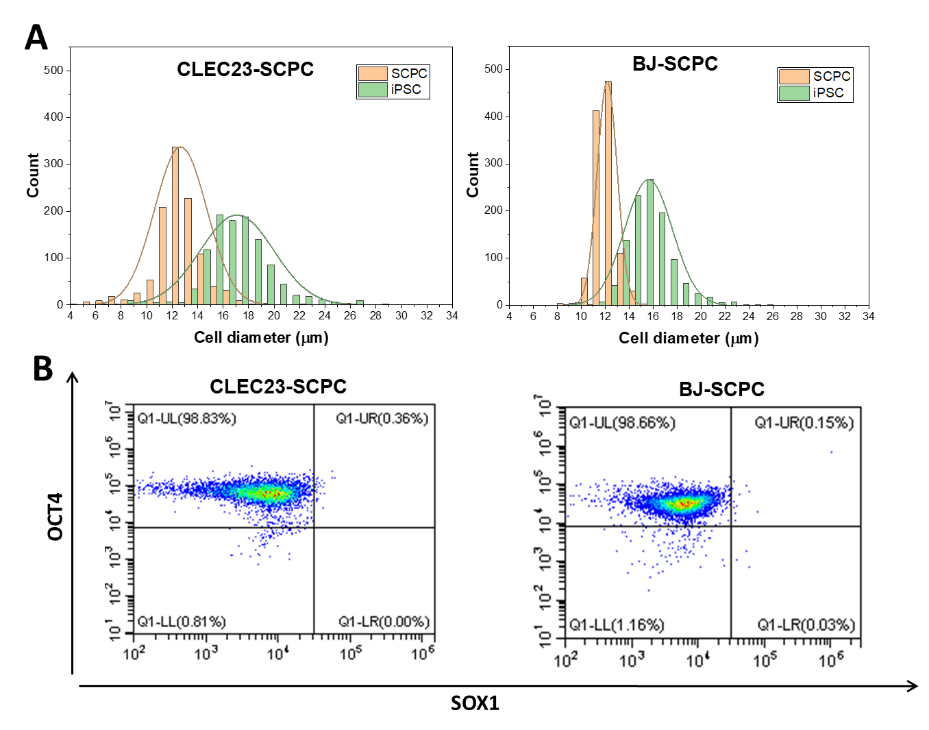

Figure S2 (A) Size profiling of the day 10 CLEC23-SCPCs and day 10 BJ-SCPCs with their derived iPSCs. It can be seen that BJ-SCPC size is more uniform than CLEC23-SCPCs. (B) Flow cytometry analysis of CLEC23-iPSCs and BJ-iPSCs with OCT4 and SOX1. Results show that >98% of cells expressed OCT4.

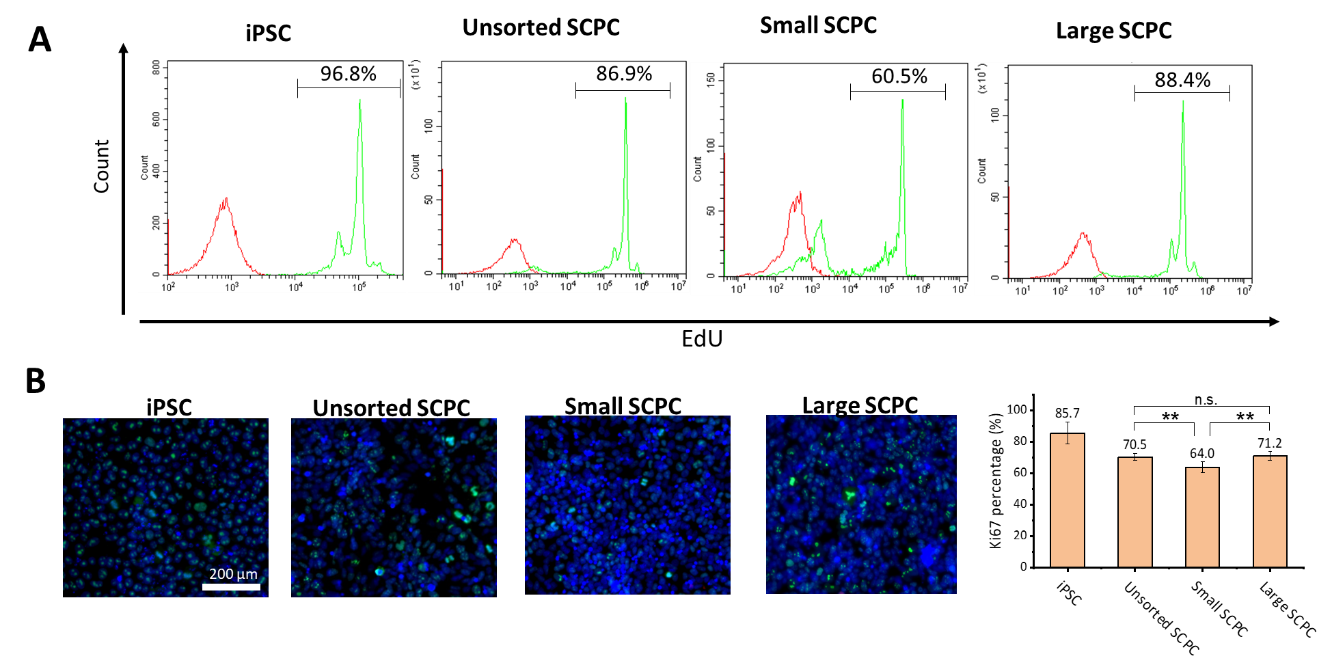

Figure S3 Proliferation assays using EdU with flow cytometry (A) and Ki67 with immunostaining analysis (B) were performed on iPSC (day 1), unsorted, and sorted (small and large) SCPCs at day 10.

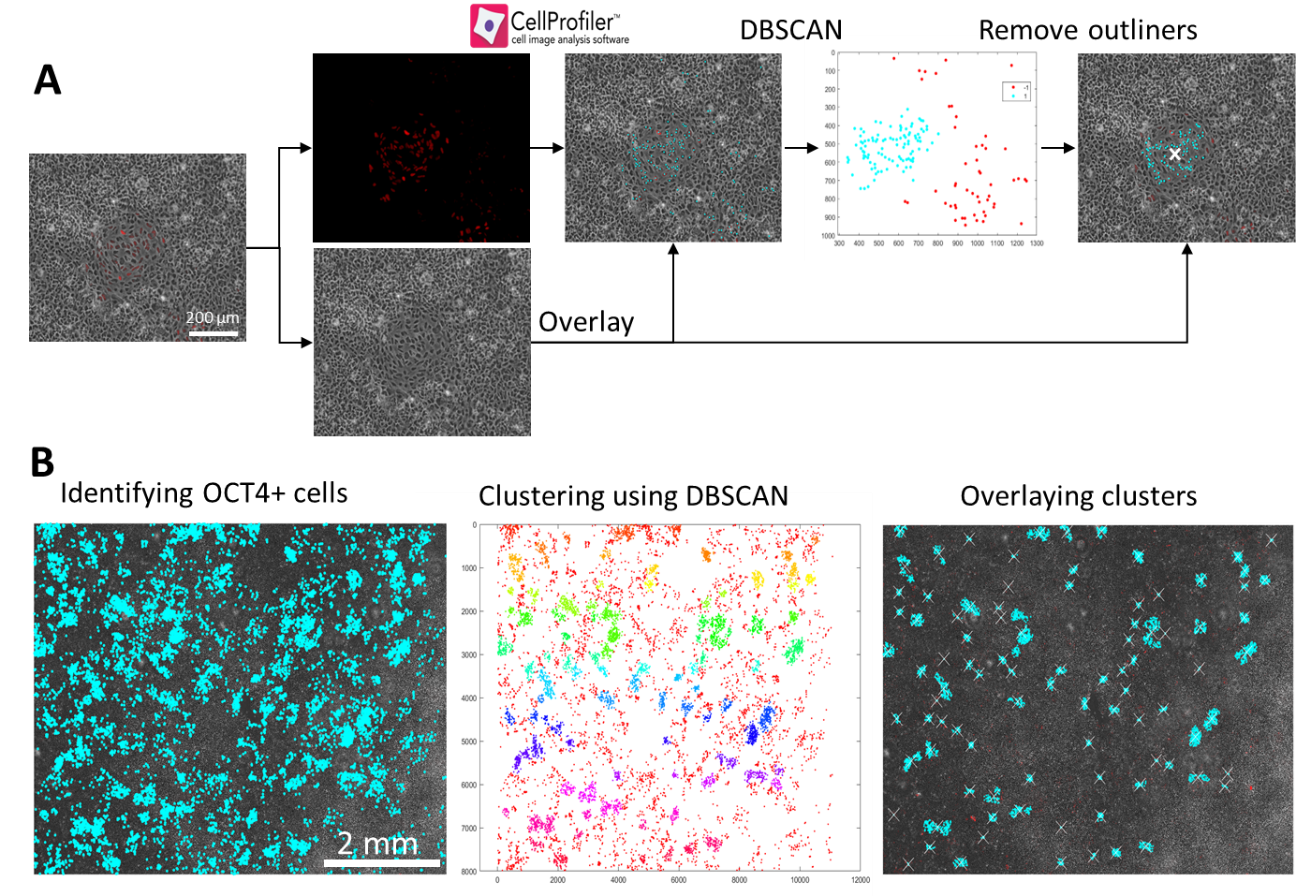

Figure S4 (A) Pipeline of the process for clustering immature colony from a single OCT4 staining image. (B) Quantified colonies colocalize with OCT4 markers in the 10 by 10 images.

Figure S4A shows the pipeline of the process for clustering immature colonies from a single OCT4 staining image. Initially, the location of OCT4+ cells in the OCT4 staining image was identified by Cell Profiler and verified by localizing with the phase contrast map. The location map of OCT4^+^ cells was then analyzed and clustered by utilizing the density-based spatial clustering of applications with noise (DBSCAN) algorithm via MATLAB. Thereby, contiguous and high-density regions of OCT4+ cells were clustered (*i.e.*, label by 1) and separated from other low point densities (*i.e.*, label by -1). The identified clusters were verified with their morphology from the phase contrast image. In order to process more colonies, this pipeline was then implemented for a map made of 10 by 10 images (100 images, Figure S4B).
